# Cell-free glycoengineering of the recombinant SARS-CoV-2 spike glycoprotein

**DOI:** 10.1101/2021.04.30.442139

**Authors:** Johannes Ruhnau, Valerian Grote, Mariana Juarez-Osorio, Dunja Bruder, Erdmann Rapp, Thomas F. T. Rexer, Udo Reichl

## Abstract

The baculovirus-insect cell expression system is readily utilized to produce viral glycoproteins for research as well as for subunit vaccines and vaccine candidates, for instance against SARS-CoV-2 infections. However, the glycoforms of recombinant proteins derived from this expression system are inherently different from mammalian cell-derived glycoforms with mainly complex-type *N*-glycans attached, and the impact of these differences in protein glycosylation on the immunogenicity is severely underinvestigated. This applies also to the SARS-CoV-2 spike glycoprotein, which is the antigen target of all licensed vaccines and vaccine candidates including virus like particles and subunit vaccines that are variants of the spike protein. Here, we expressed the transmembrane-deleted human β-1,2 N-acetlyglucosamintransferases I and II (MGAT1∆TM and MGAT2∆TM) and the β-1,4-galactosyltransferase (GalT∆TM) in *E. coli* to *in-vitro* remodel the *N*-glycans of a recombinant SARS-CoV-2 spike glycoprotein derived from insect cells. In a cell-free sequential one-pot reaction, fucosylated and afucosylated paucimannose-type *N*-glycans were converted to complex-type galactosylated *N*-glycans. In the future, this *in-vitro* glycoengineering approach can be used to efficiently generate a wide range of *N*-glycans on antigens considered as vaccine candidates for animal trials and preclinical testing to better characterize the impact of *N*-glycosylation on immunity and to improve the efficacy of protein subunit vaccines.

## 1 Introduction

Most epidemics caused by viral infections that are associated with a significant death toll were caused by enveloped viruses such as influenza A virus, human immunodeficiency virus (HIV), Zika virus, Yellow fever virus, Dengue virus and Ebolavirus. Often, the main target for neutralizing antibodies to evoke a strong immune response is a glycosylated envelope membrane protein. Thus, in the development of vaccines, glycoproteins are typically in the focus of interest. In general, the glycosylation of proteins plays a critical role regarding structure, function, solubility, stability, trafficking, and ligand-binding (Imperiali and O’Connor, 1999;Dalziel et al., 2014;Varki, 2017). Furthermore, glycosylation plays a major role for pharmacokinetics and pharmacodynamics of biologics and for pathogen-host interaction (Bagdonaite and Wandall, 2018;Cymer et al., 2018;Watanabe et al., 2019). In viral pathogenesis, glycosylation affects the attachment and release of virus particles as well as immune evasion (Schön et al.;Bagdonaite and Wandall, 2018;Watanabe et al., 2019). Especially the latter is a major hurdle for vaccine design. The mode of actions that are known to be employed to invade the immune system are secretion and shedding of glycoproteins that function as a decoy to the immune system, and the shielding of epitopes (Watanabe et al., 2019). The latter is facilitated by occluding antigenic epitopes with host-derived glycans that are obtained through hijacking the host’s cellular glycosylation machinery (Pralow et al.;Schwarzer et al., 2009;Francica et al., 2010;Helle et al., 2011;Rödig et al., 2011;Rödig et al., 2013;Sommerstein et al., 2015;Behrens et al., 2016;Gram et al., 2016;Walls et al., 2016). Moreover, it has been shown that also the glycoform itself can have an impact on binding and transmission assay as well as on transmissibility, antigenicity, and immunogenicity in animal models (Lin et al., 2003;Hütter et al., 2013;Chen et al., 2014;Li et al., 2016;Go et al., 2017). While it is assumed that immunogenic antigens benefit from mimicking the glycosylation of host cell proteins, it has also been proposed that modification of specific terminal sugar residues could be used to amplify vaccine efficacy (Galili, 2020;Chen, 2021). However, due to the complexity of protein glycosylation and the prevailing lack of methods to introduce defined modifications in the glycan composition of the proteins of interest, the topic is underinvestigated (Schön et al.;Watanabe et al., 2019;Grant et al., 2020).

The ongoing corona virus disease 2019 (COVID-19) pandemic is caused by the severe acute respiratory syndrome coronavirus 2 (SARS-CoV-2) – a single-stranded, positive-sense RNA virus (Walls et al., 2020). Its membrane envelope consists of three membrane proteins: the surface spike (S) glycoprotein, an integral membrane protein and an envelope protein (Wan et al., 2020;Zhou et al., 2020). Virus entry into human host cells is mediated by the S glycoprotein that binds to angiotensin-converting enzyme 2 (Walls et al., 2020). The S protein has 22 N-linked glycosylation sites. Thus, it is significantly more glycosylated than, for instance, the influenza A hemagglutinin (Wrapp et al., 2020). For the SARS-CoV-1 spike protein it has been shown previously that *N*-glycans significantly impact antibody response and neutralizing antibody levels (Chen et al., 2014; Walls et al., 2020).

For the investigation of the impact of glycoforms on the immunogenicity, mainly animal cell lines such as HEK and CHO cells that produce differentially glycosylated proteins are employed (Schön et al.;Lin et al., 2013). However, due to need to develop specific expression protocols for each cell line, this approach is highly work-intensive. Additionally, the inherent macro- and microheterogeneity of glycoproteins complicate the elucidation of the role of specific glycans in, for instance, regarding their immunogenicity in animal models.

Over the past years the establishment of protocols for expression of eukaryotic and bacterial glycosyltransferases has facilitated the processing of glycans in cell-free one-pot reactions. As a platform technology, the corresponding *in-vitro* glycoengineering approaches have the potential to tailor the glycoform of proteins independent of the expression systems used (Van Landuyt et al., 2019;Rexer et al., 2020a). In our study, recombinant human β-1,2 N-acetlyglucosamintransferases I and II (MGAT1∆TM and MGAT2 ∆TM) and the β-1,4-galactosyltransferase (GalT∆TM) expressed in *E. coli* were utilized to convert insect cell-derived paucimannose structures of recombinant SARS-CoV-2 spike glycoprotein to typical mammalian, complex-type galactosylated structures in a cell-free one-pot reaction (Fujiyama et al., 2001;Boeggeman et al., 2003;Bendiak, 2014;Ramakrishnan and Qasba, 2014;Stanley, 2014). Glycan structures were analyzed using multiplexed capillary gel electrophoresis with laser-induced fluorescence detection (xCGE-LIF) and Matrix-assisted laser desorption/ionization time-of-flight mass spectrometry (MALDI-TOF-MS). Results obtained clearly demonstrate that a large fraction of fucosylated and afucosylated, Man3-glycans were transferred to biantennary G2 and G2F structures (also see Table 1).

**Table 1.**
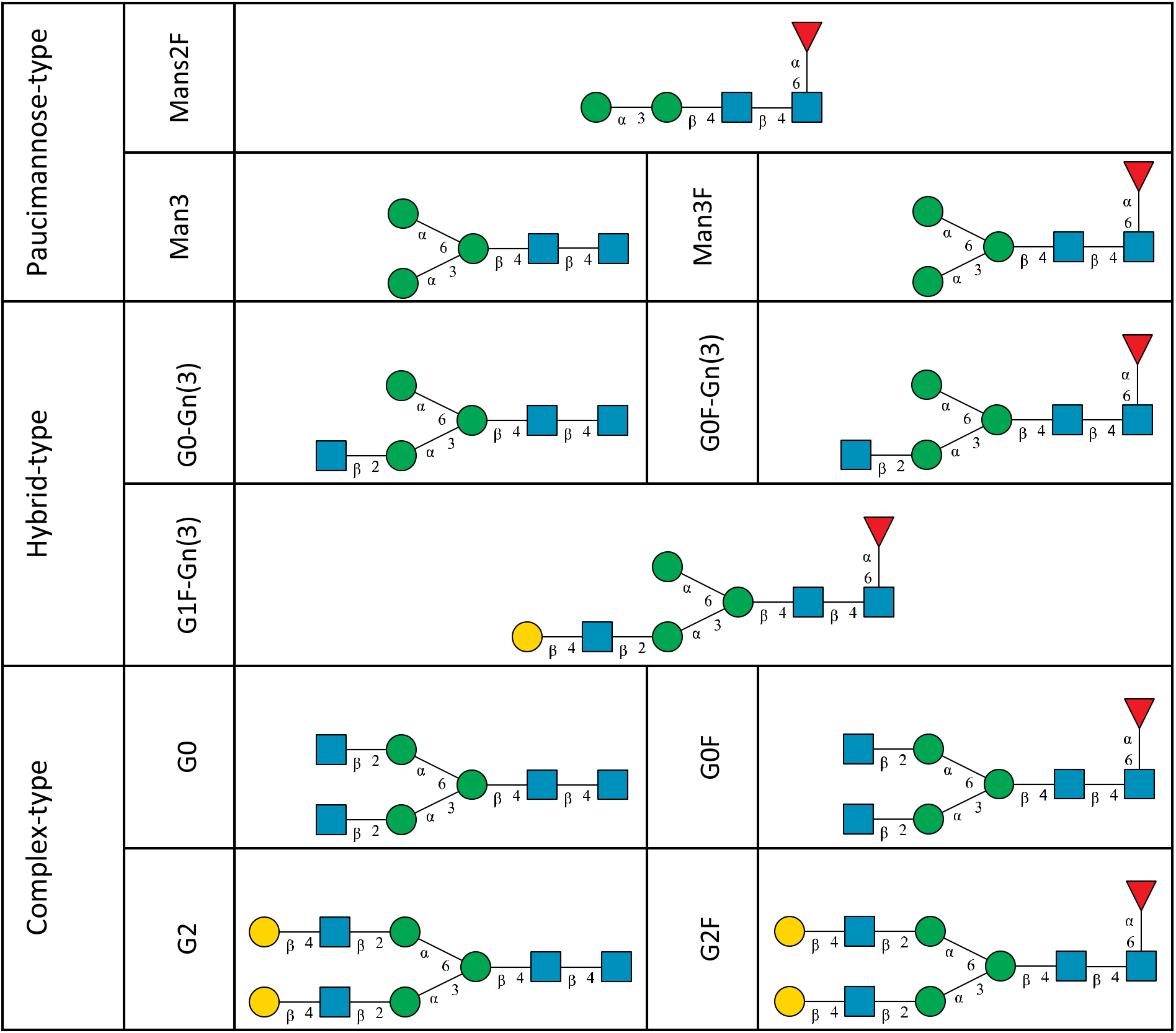
*N*-glycan categories and nomenclature for all detected and referenced structures with the exception of oligomannose-type *N*-glycans. The monosaccharide building blocks are mannose (green circle), GlcNAc (blue square), fucose (red triangle) and galactose (yellow circle).

## 2 Materials and Methods

### 2.1 Enzymes

SARS-CoV-2 spike protein containing the S1 subunit and the S2 subunit ectodomain was purchased from SinoBiologica (Beijing, PR China). The recombinant protein was produced using the baculovirus-insect-cell expression system using High-Five™; cells. The protein bears a C-terminal His-tag. For all other materials see supporting information (SI).

#### 2.1.1 Gene expression

Genes encoding for the transmembrane deleted (∆TM) variants of *Homo sapiens* α-1,3-mannosyl-glycoprotein 2-β-N-acetylglucosaminyltransferase (MGAT1∆TM) (E.C. 2.4.1.201), α-1,6-mannosyl-glycoprotein 2-β-N-acetylglucosaminyltransferase (MGAT2∆TM) (E.C. 2.4.1.143) and β-N-acetylglucosaminylglycopeptide β-1,4-galactosyltransferase (GalT∆TM) (E.C. 2.4.1.38) were expressed in *E. coli*. All constructs are bearing a 6 x histidine-tag (His-tag). For information on the cultivation, strains and vectors used see SI.

#### 2.1.2 Purification by ion metal affinity chromatography

*E. coli* cells were lysed at 4°C by high-pressure cell disruption (3 cycles, 400-600 bar) using an HPL6 homogenizer (Maximator GmbH, Nordhausen, Germany) followed by centrifugation at 7200 × g for 20 min at 4°C to precipitate cell debris. The overexpressed enzymes were filtered through 8 μm syringe filters and then purified by ion metal chromatography using an ÄKTA™; start system equipped with HisTrap™; HP columns (1 mL) (both GE Healthcare Life Sciences, Little Chalfont, UK). A buffer exchange was carried out to remove excess imidazole using an Amicon^®^ Ultra-15 Centrifugal Filter Unit – 3 kDa MW cutoff (UFC900308, Darmstadt, Germany) using standard procedures. Enzymes were stored in 50% (v/v) glycerol stock solutions at -20°C. Enzyme concentrations were determined by performing a bicinchoninic acid (BCA) assay using the Pierce™; BCA Protein Assay Kit (Thermo Fisher Scientific; Waltham, USA).

### 2.2 One-pot *in-vitro* glycoengineering reactions

Reactions were performed by sequential addition of enzymes in buffered (25 mM HEPES, pH 6.5) aqueous solutions supplemented with 10 mM MnCl_2_ at 37°C under shaking (550 rpm). The initial reaction volume (1 mL) contained 0.1 μg/mL of SARS-CoV-2 spike protein, 4 mM UDP-GlcNAc and 0.2 μg/μL MGAT1∆TM. After a reaction time of 12 h, 150 μL of a buffered solution containing 4 mM UDP-GlcNAc and 0.85 μg/μL MGAT2∆TM was added to 500 μL of the reaction. After 12 more hours, 175 μL of a buffered solution containing 4 mM UDP-galactose and 0.56 μg/μL GalT∆TM was added to 325 μL of the reaction mix. Aliquots of the reactions were taken for *N*-glycan analysis before the addition of each enzyme and at the end of the reaction (12 h after GalT∆TM addition).

#### 2.2.1 Sample pretreatment: PNGase F digest of N-glycosylated proteins

1 μg *N*-glycosylated protein sample was linearized and denatured by adding 2 μL 2 % (w/v) SDS in PBS buffer (pH 7.2) and subsequent heating at 60°C for 10 min. Samples were cooled down to room temperature. 4 μL 8 % (w/v) IGEPAL in PBS and 1 μL of a 1 U/μL PNGase F solution were added. Samples were incubated for 1 h at 37°C, vacuum evaporated and dissolved in 20 μL LC-MS grade H_2_O.

#### 2.2.2 xCGE-LIF-based N-Glycan Analysis

*N*-glycan analysis based on xCGE-LIF was conducted using a glyXboxCE™;-system (glyXera, Magdeburg, Germany) according to (Hennig et al., 2015;Hennig et al., 2016). Briefly, 2 μL of each sample was used for fluorescent labelling of *N*-glycans with 8-aminopyrene-1,3,6-trisulfonic acid (APTS) following post derivatization clean-up by hydrophilic interaction liquid chromatography-solid phase extraction (HILIC-SPE) with the glyXprep16™; kit (glyXera). Data processing, normalization of migration times and annotation of *N*-glycan fingerprints were performed with glyXtool™; software (glyXera).

#### 2.2.3 MALDI-TOF-MS-based N-Glycan Analysis

MALDI-TOF-MS analysis of released *N*-glycans was performed as described previously (Selman et al., 2011;Fischöder et al., 2019). Briefly, 0.9 cm cotton rope was used for Cotton HILIC SPE. The stationary phase was equilibrated with 50 μL LC-MS grade H_2_O followed by 50 μL 85% ACN_aq_. 10 μL of released *N*-glycans were adjusted to 70 μL 85% ACN_aq_ with 1% TFA and loaded onto the HILIC phase. Following 2 washing steps with 50 μL 85% ACN_aq_ with 1% TFA and 50 μL 85% ACN_aq_, the samples were eluted in 70 μL LC-MS grade H_2_O, vacuum evaporated and dissolved in 20 μL LC-MS grade H_2_O. For the MALDI-TOF-MS analysis 0.5 μL super-dihydroxybenzoic acid (S-DHB) (≥99.0%, Sigma-Aldrich, Steinheim, Germany) matrix (10 mg/mL) in 30% (v/v) ACN_aq_, 0.1% (v/v) TFA, 2 mM NaCl was spotted onto a MTP AnchorChip 800/384 TF MALDI target (Bruker Daltonics, Bremen, Germany). Subsequently 1 μL sample was applied onto the dried matrix layer. Measurements were carried out on an ultrafleXtreme MALDI-TOF/TOF MS (Bruker Daltonics, Bremen, Germany) in reflectron positive ion mode. Data was processed with the top-hat filter and the adjacent-averaging algorithm using flexAnalysis version 3.3 Build 80 (Bruker Daltonics, Bremen, Germany).

### 2.3 *N*-Glycan Nomenclature

*N*-Glycan nomenclature was adopted from Stanley et al (Stanley et al., 2015). Depiction of *N*-glycan structures followed the Symbol Nomenclature for Glycans (SNFG) guidelines (Neelamegham et al., 2019). The *N*-glycan sketches in this manuscript were produced using the “Glycan Builder2” software tool (Tsuchiya et al., 2017). *N*-Glycans are typically categorized into paucimannose-, oligomannose-, hybrid- and complex-type structures.

## 3 Results

### 3.1 Pathway design

The human *in-vivo* cascade reaction for the generation of complex-type *N*-glycans from the conserved ER-derived oligomannose-type *N*-glycan precursor GlcNAc_2_Man_9_Glc_3_, was in part remodelled *in-vitro* to generate fully galactosylated complex-type *N*-glycans starting from insect cell-derived paucimannose-type *N*-glycans. Central to the construction of the simplified *in-vitro* cascade is the ability of human MGAT1 to utilize Man3 and Man3F as substrates, which allows circumventing the application of recombinant mannosidases (see Figure 1). For the production of the G2 structure from paucimannose-type *N*-glycans, the three recombinant glycosyltransferases MGAT1∆TM, MGAT2∆TM and GalT∆TM were successfully produced in *E. coli* (see SI). Enzyme concentrations of typically 1.3 mg/mL after ion metal affinity chromatography (IMAC) and buffer exchange were obtained. In scouting experiments, it was confirmed that all enzymes are active in the buffered solutions (pH 6.5) with MnCl_2_ supplemented as a co-factor (data not shown).

**Figure 1.**
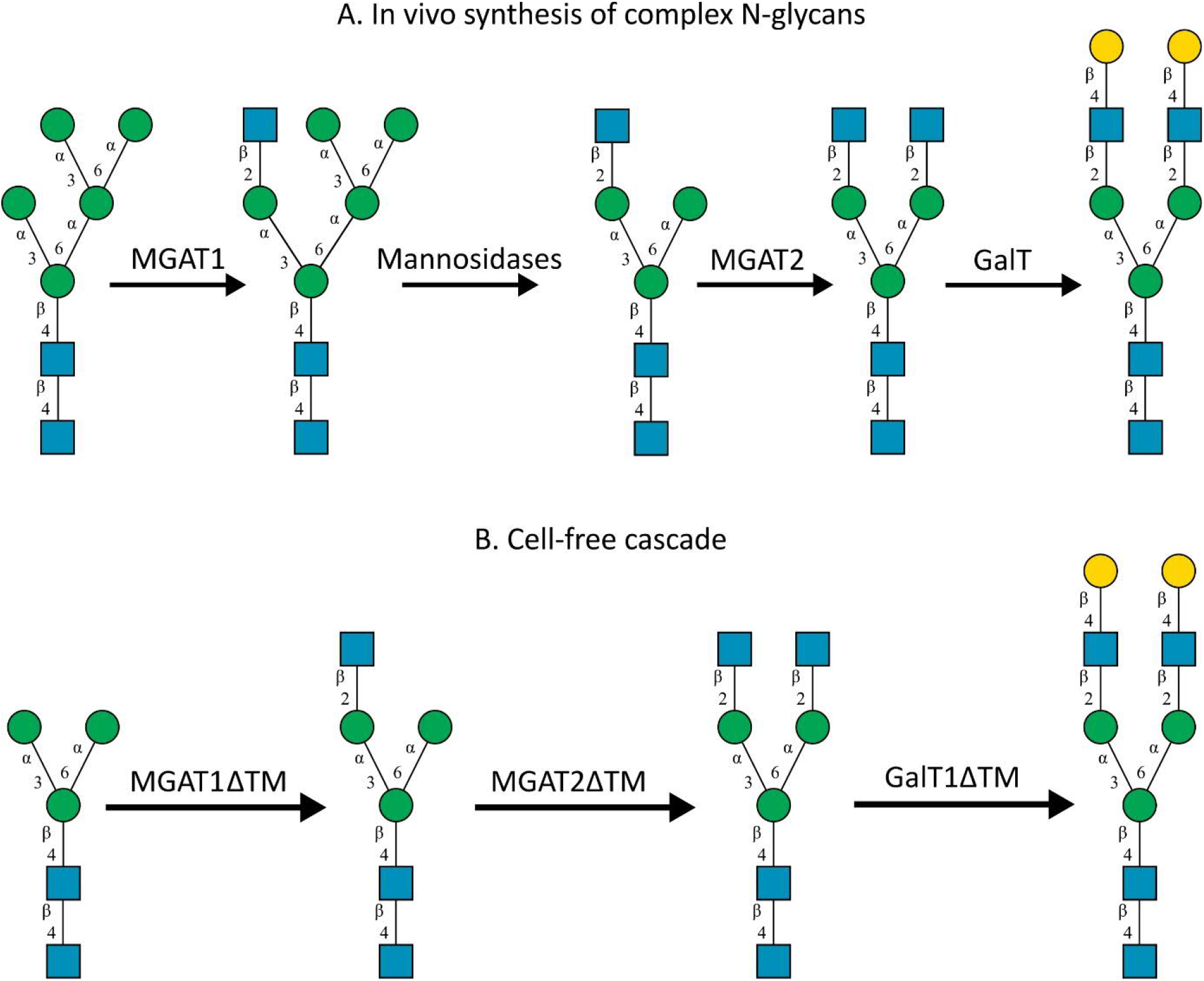
A) *In-vivo* the oligomannose-type *N*-glycan Man5 is converted into complex-type *N*-glycans by mannosidases and MGAT1, MGAT2 and GalT. Substrates for these reactions are UDP-GlcNAc and UDP-galactose, respectively. B) This process can be remodelled *in-vitro* to synthesize complex-type structures on insect cell-derived recombinant proteins with paucimannose-type N-glycans, like Man3.

### 3.2 Glycoform of the unprocessed recombinant SARS-CoV-2 spike glycoprotein

Analytical characterization of the unprocessed and glycoengineered SARS-CoV 2 spike protein was achieved by the two orthogonal methods xCGE-LIF and MALDI-TOF-MS (see Figure 2 and Figure 3). The high-resolution *N*-glycan fingerprints (migration time aligned & and peak height normalized electropherograms) from xCGE-LIF combined with the precise mass profiles generated by MALDI-TOF-MS allowed for fast and robust annotation also of isomeric *N*-glycan structures. Furthermore, normalization of *N*-glycan fingerprints to total peak height enabled relative quantification of individual *N*-glycan structures by xCGE-LIF. The insect-cell-produced recombinant SARS-CoV-2 spike glycoprotein prominently displays α-1,6-core-fucosylated Man3F and G0F-Gn(3) structures (Figure 2A blue and Figure 3A). Moreover, Man2F, Man3, the hybrid-type structure G0-Gn(3), the complex-type structure G0F, and afucosylated oligomannose-type structures were detected. There is excellent agreement between xCGE-LIF and MALDI-TOF-MS measurements.

**Figure 2.**
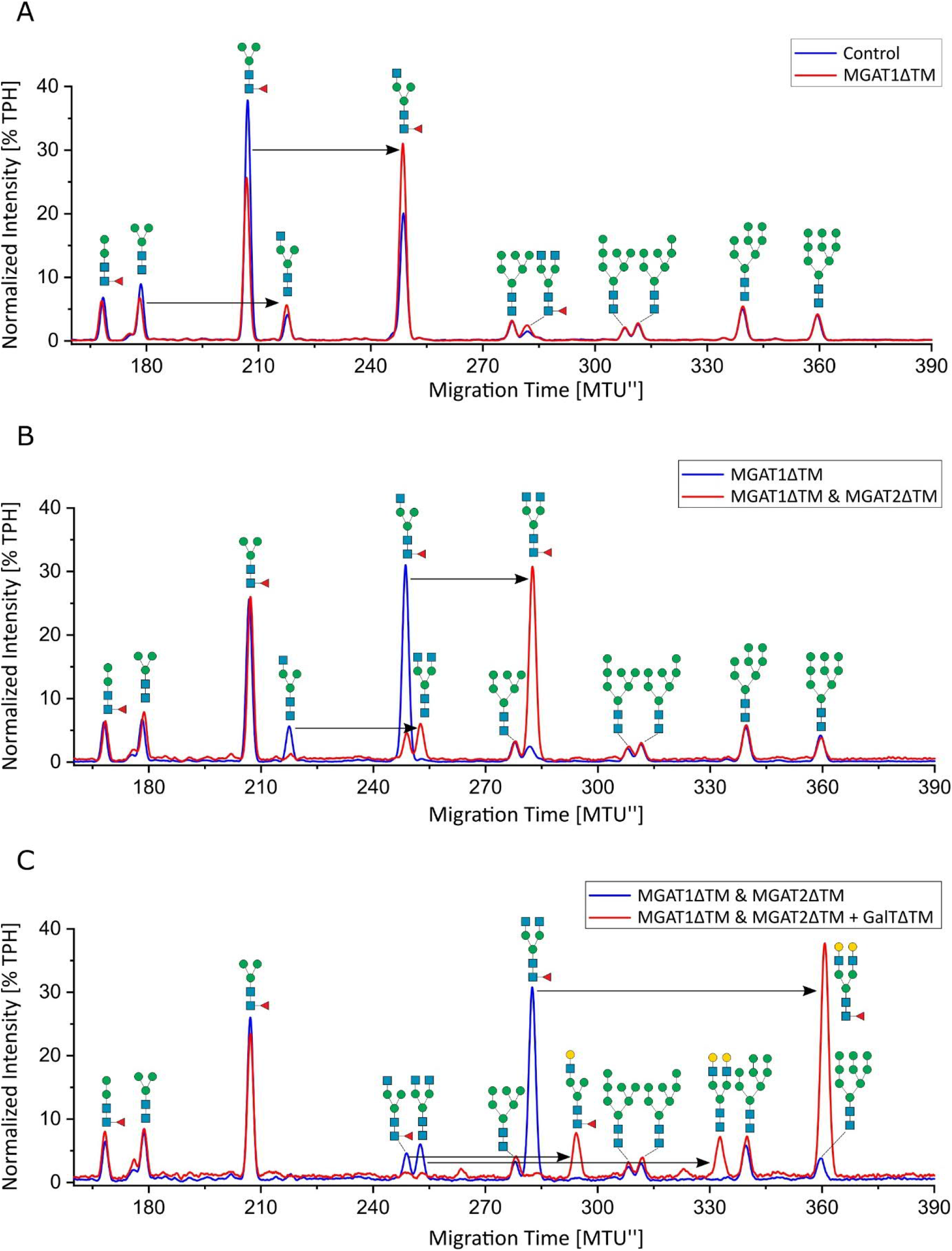
xCGE-LIF *N*-glycan fingerprints of unprocessed and *in-vitro* glycoengineered SARS-CoV-2 spike protein *N*-glycans. *N*-glycosylation pattern of the spike protein: A) unprocessed (blue) and 12 h after start of the reaction with MGAT1∆TM (red); B) 12 h after start of the reaction with MGAT1∆TM (blue) and 12 h after the addition of MGAT2∆TM (red); C) 12 h after the addition of MGAT2∆TM (blue) and 12 h after addition of GalT∆TM (red). TPH, total peak height.

**Figure 3.**
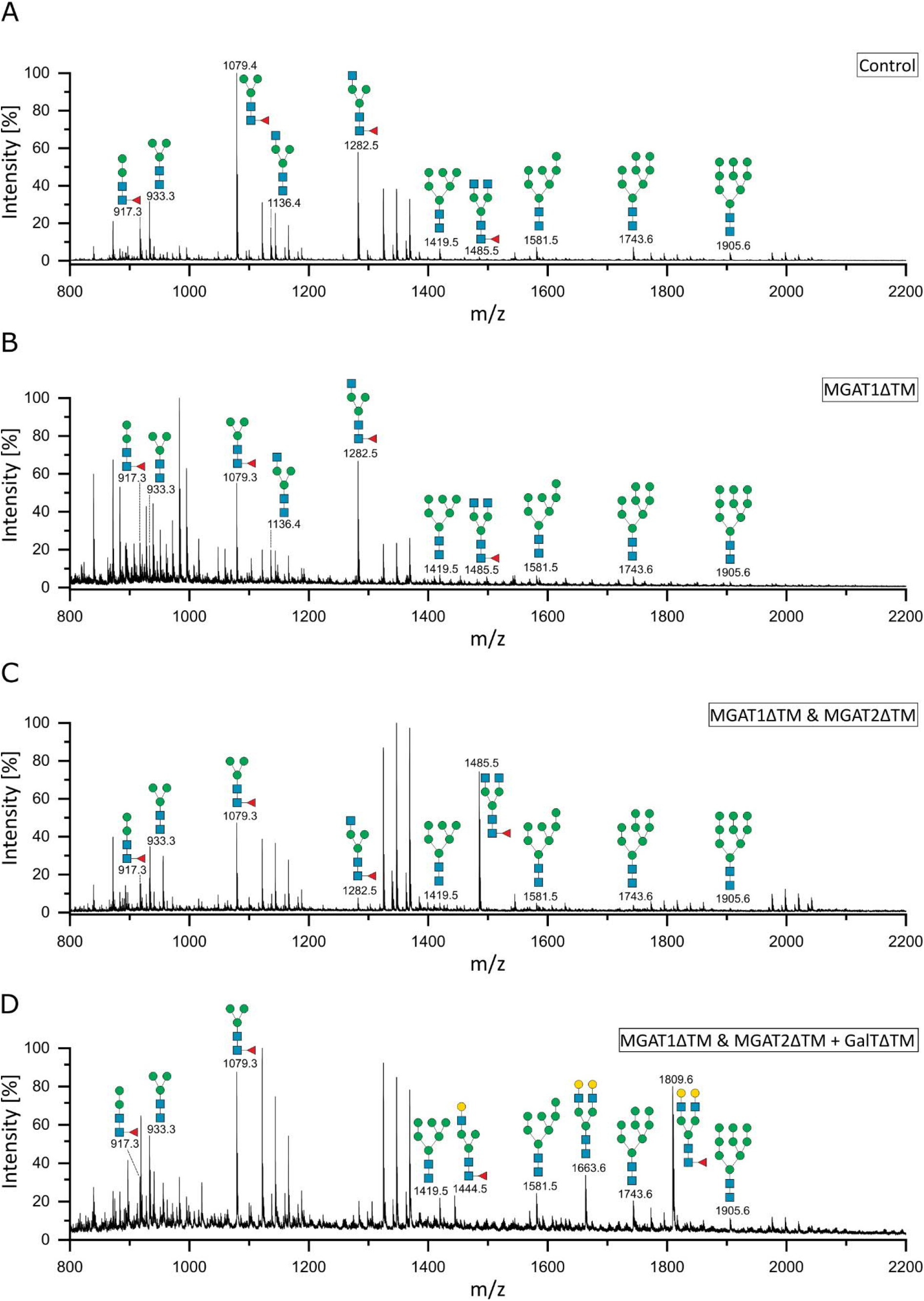
MALDI-TOF mass spectra of the unprocessed and glycoengineered SARS-COV 2 spike protein glycoforms. *N*-glycans were detected in reflectron positive ion mode as sodium adducts ([M+Na]+). A) Unprocessed spike protein. B.) 12 h after start of the reaction with MGAT1∆TM. C) 12 h after the addition of MGAT2∆TM. D) 12 h after the addition of GalT∆TM. Only the peaks that depict *N*-glycans are annotated.

### 3.3 *In-vitro* glycoengineering of SARS-CoV-2 spike glycoprotein

Recombinant MGAT1∆TM, MGAT2∆TM and GalT∆TM were used in a one-pot glycoengineering reaction to convert the paucimannose structures to complex-type *N*-glycans. In scouting experiments it was found that after MGAT1∆TM, MGAT2∆TM and GalT∆TM addition at the start of the reaction, Man3F was converted to, at least in parts, to the hybrid-type structure G1F-Gn(3) missing the extension on the α1-6 mannosylated antenna catalysed by MGAT2. G1F-Gn(3) is not a natural substrate for MGAT2 and can, if at all, most likely only be processed at very low turnover rates. Thus, the reactions were carried out by adding the enzymes sequentially as detailed in M&M. In the first step, a GlcNAc residue is added from UDP-GlcNAc to the α-1,3-linked terminal mannose antenna of Man3F and Man3 by MGAT1∆TM (see Figure 2A and Figure3A and B). After a reaction time of 12 h only parts of Man3 and Man3F were converted to G0-Gn(3) and G0F-Gn(3), respectively. Scouting experiment showed that the conversion is typically irreversible and, thus, the incomplete processing is either due to low turnover or possible enzyme inactivation of MGAT1∆TM. Another possibility is that the glycans are inaccessible for MGAT1∆TM but can be released from the backbone by PNGase F. In the second step, UDP-GlcNAc and MGAT2∆TM are added. After incubation for 12 h, the hybrid-type structures G0-Gn(3) and G0F-Gn(3) were converted to G0 and G0F, respectively, with only a minor fraction of G0F-Gn(3) remaining (Figure 2B and Figure C). MGAT2∆TM did not show any activity towards Man3 and Man3F. However, as mentioned before, this could be due the inaccessibility of these glycans. In the final step, the reaction was supplemented with UDP-galactose and GalT∆TM to add galactose to the terminal GlcNAc. At the end point of the reaction, after another 12 h of incubation, the galactosylated complex-type structure G2 was dominant along unprocessed Man3F (see Figure 2C and Figure 3D). Moreover, G0 was completely converted to G2 while the residual amount of the hybrid-type structure G0F-Gn(3) was also galactosylated to G1F-Gn(3). All oligomannose-type structures remained unaltered throughout the reaction. In general, the xCGE-LIF and the MALDI-TOF-MS data were in excellent agreement for all measurements.

## 4 Discussion

Due to its scalability, eukaryotic protein processing and high productivity, the baculovirus-insect cell expression system is well-suited for the production of subunit vaccines (Palomares et al.;Felberbaum, 2015). In addition to subunit vaccines against SARS-CoV-2 infections in development, there are currently three licensed vaccines, Flublok^®^, Cervarix^®^ and Provenge^®^ produced using this expression system with several more in clinical trials (Palomares et al.;Felberbaum, 2015).

High immunogenicity of a recombinant insect-cell produced spike protein ectodomain variant, very similar to the one used here, has been confirmed in non-human primates [40]. Moreover, the spike protein is the antigen target of virtually all COVID-19 vaccines and advanced vaccine candidates (Krammer, 2020). At the time of writing this article, there was one licensed COVID-19 protein subunit vaccine (RBD-Dimer from Anhui Zhifei Longcom Biopharmaceutical, China) in China, while for two more candidates (Covovax from Novavax, USA; VAT00002 from Sanofi Pasteur and GSK, France / UK) emergency authorization was pending in the US and Europe (Yang et al., 2020;Dai and Gao, 2021;Shrotri et al., 2021). All three are recombinant SARS-CoV-2 spike protein variants produced using the baculovirus-insect cell expression system (Kyriakidis et al., 2021).

Glycoforms of recombinant proteins produced using baculovirus-insect cell expression systems are profoundly different from those produced using mammalian expression systems. An extensive review on the glycosylation processing of insect cells is given by Geisler et al (Geisler et al., 2015). Typically, these proteins display mainly paucimannose and hybrid-type *N*-glycans with, at most, minor fractions of complex-type and oligomannose-type *N*-glycans (Palomares et al.;Geisler et al., 2015). In this respect, the glycoform we observed on the pure spike protein variants is exemplary for an insect cell protein expression.

For vaccine development, it has been proposed that immunogen candidates benefit from closely mimicking the macro- and microheterogeneity of the live virus glycosylation (Grant et al., 2020;Watanabe et al., 2020). This is as eliciting antibodies against shielded or non-native epitopes could cause an inefficient immune response. To overcome such obstacles, novel strategies utilizing distinct non-human glycans containing N-glycolylneuraminic acid or α,1-3 linked galactose residues, have been proposed to alleviate immune responses (Schön et al.;Hütter et al., 2013;Galili, 2020;Chen, 2021). However, such approaches still need to be investigated in detail experimentally as, for example, both compounds are also suspected to cause allergenic reactions in humans.

To convert the glycoform from primarily paucimannose-type to typical mammalian complex-type *N*-glycans, the recombinant human glycosyltransferases, MGAT1∆TM, MGAT2∆TM and GalT∆TM, were effectively combined in a cell-free, one-pot glycosylation reaction. The gene expression of these glycosyltransferases in *E. coli* and the activity of the His-tag purified, soluble recombinant proteins in one-pot reactions using free glycans as substrates has been shown before ((Fujiyama et al., 2001), Mahour *et al.*, unpublished).

The site-specific glycan analysis of recombinant SARS-CoV-2 spike protein ectodomain expressed in human-derived cell line FreeStyle™; 293-F showed that of the 22 *N*-glycosylation sites only eight contained substantial fractions of oligomannose-type *N*-glycans (Watanabe et al., 2020). It is assumed that the occurrence of oligomannose-type fractions is caused by the steric inaccessibility of these glycans to the glycan processing enzymes in the Golgi, i.e. the occurrence of oligomannose-type *N*-glycans at distinct sites has shown to be independent of the producer cell line for the HIV viral glycoprotein gp120 (Pritchard et al., 2015). In accordance with the human cell-derived spike protein, our engineered spike protein abundantly exhibited complex-type G2F *N*-glycans. To a minor extend, a range of hybrid- and oligomannose-type *N*-glycans were also detected on the engineered spike protein. In contrast to the engineered spike protein, human cell-derived spike proteins also exhibit complex-type multi-antennary and sialylated structures (Watanabe et al., 2020). Taken together, a significant overlap of the glycoform has been generated. Whether the overlap is also site-specific remains to be investigated in future.

Over the past years, many efforts have been made to engineer insect cell lines to express complex-type *N*-glycans. A comprehensive summary of the attempts is given by Palomares et al. (Palomares et al.). Briefly, complex-type *N*-glycans can be produced by the co-expression of glycosyltransferases or by generating transient insect cell lines. While the former generates an additional metabolic burden and affects growth properties, the stability of the latter has not been examined for commercial scale use. The advantage of *in-vitro* glycoengineering lies in its independence of producer cell lines as well as its flexibility towards the option to readily generate different glycoforms that are close to homogeneity. However, expensive nucleotides sugars are required as substrates and, thus, it is so far not feasible to apply *in-vitro* glycoengineering at larger scales (Mahour et al., 2018;Rexer et al., 2018;Rexer et al., 2020b).

## 5 Conclusion

SARS-CoV-2 spike glycoprotein variants produced in a baculovirus-insect cell expression system were *in-vitro* glycoengineered using recombinant glycosyltransferases to mimic the glycoform observed on the human cell-derived protein. *In-vitro* glycoengineering reactions as conducted here, can be used to generate immunogen candidates for pre-clinical testing to investigate the role of glycosylation on the antigenicity and immunogenicity in animal models. In general, *in-vitro* glycoengineering approaches can virtually be used to tailor the glycoform of all prominent vaccine candidates such as activated and attenuated viruses and virus like particles. The application of the technology to larger scales depends on the bulk availability of sugar nucleotides at moderate costs.

## 6 Acknowledgement

The project is funded by the Deutsche Forschungsgemeinschaft (DFG, German Research Foundation) – Projektnummer 458633485. Dunja Bruder acknowledges funding from the State of Saxony-Anhalt (Förderkennzeichen I 130). Erdmann Rapp und Valerian Grote acknowledge funding by DFG-FOR2509: RA2992/1-1. Furthermore, the authors would like to thank Anja Bastian for her excellent technical support. Lisa Wenzel and Reza Mahour are acknowledged for help with the expression and purification of glycosyltransferases.

---

**8.** List of abbreviations.

8. List of abbreviations
|  |  |
| --- | --- |
| APTS | 8-Aminopyrene-1,3,6-trisulfonic acid |
| dH<sub>2</sub>O | Deionized water |
| GlcNAc | *N*-acetylglucosamine |
| HEPES | 4-(2-hydroxyethyl)-1-piperazineethanesulfonic acid |
| HILIC | Hydrophilic interaction liquid chromatography |
| His tag | Histidine tag |
| IMAC | Immobilized Metal Affinity Chromatography |
| IPTG | Isopropyl β-D-1-thiogalactopyranoside |
| LC | Liquid Chromatography |
| MALDI-TOF | Matrix-assisted laser desorption/ionization time-of-flight mass spectrometry |
| MS | Mass spectrometry |
| MnCl<sub>2</sub> | Mangan(II)-chlorid |
| MTU'' | Migration Time Units after alignment to internal standards |
| rpm | Rounds per min |
| SDS-PAGE | Sodium dodecyl sulfate polyacrylamide gel electrophoresis |
| SPE | Solid-phase extraction |
| TPH | Total peak height |
| UDP-galactose | Uridine-diphosphate galactose |
| UDP-GlcNAc | Uridine diphosphate *N*-acetylglucosamine |
| xCGE-LIF | Multiplexed capillary gel electrophoresis with laser-induced fluorescence detection |

## Literature

Bagdonaite, I., and Wandall, H.H. (2018). Global aspects of viral glycosylation. Glycobiology 28, 443–467.

Behrens, A.-J., Vasiljevic, S., Pritchard, L.K., Harvey, D.J., Andev, R.S., Krumm, S.A., Struwe, W.B., Cupo, A., Kumar, A., Zitzmann, N., Seabright, G.E., Kramer, H.B., Spencer, D.I.R., Royle, L., Lee, J.H., Klasse, P.J., Burton, D.R., Wilson, I.A., Ward, A.B., Sanders, R.W., Moore, J.P., Doores, K.J., and Crispin, M. (2016). Composition and Antigenic Effects of Individual Glycan Sites of a Trimeric HIV-1 Envelope Glycoprotein. Cell Reports 14, 2695–2706.

Bendiak, B. (2014). “Mannosyl (Alpha-1,6-)-Glycoprotein Beta-1,2-N-Acetylglucosaminyltransferase (MGAT2),” in Handbook of Glycosyltransferases and Related Genes, eds. N. Taniguchi, K. Honke, M. Fukuda, H. Narimatsu, Y. Yamaguchi & T. Angata. (Tokyo: Springer Japan), 195–207.

Boeggeman, E.E., Ramakrishnan, B., and Qasba, P.K. (2003). The N-terminal stem region of bovine and human β1,4-galactosyltransferase I increases the in vitro folding efficiency of their catalytic domain from inclusion bodies. Protein Expression and Purification 30, 219–229.

Chen, J.-M. (2021). SARS-CoV-2 replicating in nonprimate mammalian cells probably have critical advantages for COVID-19 vaccines due to anti-Gal antibodies: A minireview and proposals. Journal of Medical Virology 93, 351–356.

Chen, W.-H., Du, L., Chag, S.M., Ma, C., Tricoche, N., Tao, X., Seid, C.A., Hudspeth, E.M., Lustigman, S., Tseng, C.-T.K., Bottazzi, M.E., Hotez, P.J., Zhan, B., and Jiang, S. (2014). Yeast-expressed recombinant protein of the receptor-binding domain in SARS-CoV spike protein with deglycosylated forms as a SARS vaccine candidate. Human Vaccines & Immunotherapeutics 10, 648–658.

Cymer, F., Beck, H., Rohde, A., and Reusch, D. (2018). Therapeutic monoclonal antibody N-glycosylation – Structure, function and therapeutic potential. Biologicals 52, 1–11.

Dai, L., and Gao, G.F. (2021). Viral targets for vaccines against COVID-19. Nature reviews. Immunology 21, 73–82.

Dalziel, M., Crispin, M., Scanlan, C.N., Zitzmann, N., and Dwek, R.A. (2014). Emerging Principles for the Therapeutic Exploitation of Glycosylation. Science 343, 1235681–1235681-1235681–1235688.

Felberbaum, R.S. (2015). The baculovirus expression vector system: A commercial manufacturing platform for viral vaccines and gene therapy vectors. Biotechnology Journal 10, 702–714.

Fischöder, T., Cajic, S., Grote, V., Heinzler, R., Reichl, U., Franzreb, M., Rapp, E., and Elling, L. 2019. Enzymatic Cascades for Tailored ^13^C_6_ and ^15^N Enriched Human Milk Oligosaccharides. Molecules [Online], 24. Available: https://www.mdpi.com/1420-3049/24/19/3482.

Francica, J.R., Varela-Rohena, A., Medvec, A., Plesa, G., Riley, J.L., and Bates, P. 2010. Steric shielding of surface epitopes and impaired immune recognition induced by the ebola virus glycoprotein. PLoS Pathog [Online], 6. [Accessed Sep 9].

Fujiyama, K., Ido, Y., Misaki, R., Moran, D.G., Yanagihara, I., Honda, T., Nishimura, S.I., Yoshida, T., and Seki, T. (2001). Human N-Acetylglucosaminyltransferase I. Expression in Escherichia coli as a soluble enzyme, and application as an immobilized enzyme for the chemoenzymatic synthesis of N-linked oligosaccharides. Journal of Bioscience and Bioengineering 92, 569–574.

Galili, U. (2020). Amplifying immunogenicity of prospective Covid-19 vaccines by glycoengineering the coronavirus glycan-shield to present α-gal epitopes. Vaccine 38, 6487–6499.

Geisler, C., Mabashi-Asazuma, H., and Jarvis, D.L. (2015). “An overview and history of glyco-engineering in insect expression systems,” in Glyco-Engineering: Methods and Protocols.), 131–152.

Go, E.P., Ding, H., Zhang, S., Ringe, R.P., Nicely, N., Hua, D., Steinbock, R.T., Golabek, M., Alin, J., Alam, S.M., Cupo, A., Haynes, B.F., Kappes, J.C., Moore, J.P., Sodroski, J.G., and Desaire, H. 2017. Glycosylation Benchmark Profile for HIV-1 Envelope Glycoprotein Production Based on Eleven Env Trimers. Journal of Virology [Online], 91. Available: https://jvi.asm.org/content/jvi/91/9/e02428-16.full.pdf.

Gram, A.M., Oosenbrug, T., Lindenbergh, M.F.S., Bull, C., Comvalius, A., Dickson, K.J.I., Wiegant, J., Vrolijk, H., Lebbink, R.J., Wolterbeek, R., Adema, G.J., Griffioen, M., Heemskerk, M.H.M., Tscharke, D.C., Hutt-Fletcher, L.M., Wiertz, E.J.H.J., Hoeben, R.C., and Ressing, M.E. (2016). The Epstein-Barr Virus Glycoprotein gp150 Forms an Immune-Evasive Glycan Shield at the Surface of Infected Cells. Plos Pathogens 12.

Grant, O.C., Montgomery, D., Ito, K., and Woods, R.J. 2020. Analysis of the SARS-CoV-2 spike protein glycan shield reveals implications for immune recognition. Scientific Reports [Online], 10. Available: https://doi.org/10.1038/s41598-020-71748-7 [Accessed 2020/09/14].

Helle, F., Duverlie, G., and Dubuisson, J. (2011). The Hepatitis C Virus Glycan Shield and Evasion of the Humoral Immune Response. Viruses-Basel 3, 1909–1932.

Hennig, R., Cajic, S., Borowiak, M., Hoffmann, M., Kottler, R., Reichl, U., and Rapp, E. (2016). Towards personalized diagnostics via longitudinal study of the human plasma N-glycome. Biochimica et Biophysica Acta (BBA) - General Subjects 1860, 1728–1738.

Hennig, R., Rapp, E., Kottler, R., Cajic, S., Borowiak, M., and Reichl, U. (2015). “N-Glycosylation fingerprinting of viral glycoproteins by xCGE-LIF”, in: Methods in Molecular Biology. Humana Press Inc.).

Hütter, J., Rödig, J.V., Höper, D., Seeberger, P.H., Reichl, U., Rapp, E., and Lepenies, B. (2013). Toward Animal Cell Culture-based Influenza Vaccine Design: Viral Hemagglutinin N-Glycosylation Markedly Impacts Immunogenicity. The Journal of Immunology 190, 220–230.

Imperiali, B., and O’connor, S.E. (1999). Effect of N-linked glycosylation on glycopeptide and glycoprotein structure. Current Opinion in Chemical Biology 3, 643–649.

Krammer, F. (2020). SARS-CoV-2 vaccines in development. Nature 586, 516–527.

Kyriakidis, N.C., López-Cortés, A., González, E.V., Grimaldos, A.B., and Prado, E.O. 2021. SARS-CoV-2 vaccines strategies: a comprehensive review of phase 3 candidates. npj Vaccines [Online], 6. Available: https://doi.org/10.1038/s41541-021-00292-w [Accessed 2021/02/22].

Li, D., Von Schaewen, M., Wang, X., Tao, W., Zhang, Y., Li, L., Heller, B., Hrebikova, G., Deng, Q., Ploss, A., Zhong, J., and Huang, Z. (2016). Altered Glycosylation Patterns Increase Immunogenicity of a Subunit Hepatitis C Virus Vaccine, Inducing Neutralizing Antibodies Which Confer Protection in Mice. Journal of Virology 90, 10486–10498.

Lin, G., Simmons, G., Pöhlmann, S., Baribaud, F., Ni, H., Leslie, G.J., Haggarty, B.S., Bates, P., Weissman, D., Hoxie, J.A., and Doms, R.W. (2003). Differential N-Linked Glycosylation of Human Immunodeficiency Virus and Ebola Virus Envelope Glycoproteins Modulates Interactions with DC-SIGN and DC-SIGNR. Journal of Virology 77, 1337–1346.

Lin, S.-C., Jan, J.-T., Dionne, B., Butler, M., Huang, M.-H., Wu, C.-Y., Wong, C.-H., and Wu, S.-C. 2013. Different Immunity Elicited by Recombinant H5N1 Hemagglutinin Proteins Containing Pauci-Mannose, High-Mannose, or Complex Type N-Glycans. PLOS ONE [Online], 8. Available: https://doi.org/10.1371/journal.pone.0066719.

Mahour, R., Klapproth, J., Rexer, T.F.T., Schildbach, A., Klamt, S., Pietzsch, M., Rapp, E., and Reichl, U. (2018). Establishment of a five-enzyme cell-free cascade for the synthesis of uridine diphosphate N-acetylglucosamine. Journal of Biotechnology 283, 120–129.

Neelamegham, S., Aoki-Kinoshita, K., Bolton, E., Frank, M., Lisacek, F., Lütteke, T., O’boyle, N., Packer, N.H., Stanley, P., Toukach, P., Varki, A., Woods, R.J., and Group, T.S.D. (2019). Updates to the Symbol Nomenclature for Glycans guidelines. Glycobiology 29, 620–624.

Palomares, L.A., Srivastava, I.K., Ramírez, O.T., and Cox, M.M.J. “Glycobiotechnology of the Insect Cell-Baculovirus Expression System Technology.” (Berlin, Heidelberg: Springer Berlin Heidelberg), 1–22.

Pralow, A., Hoffmann, M., Nguyen-Khuong, T., Pioch, M., Hennig, R., Genzel, Y., Rapp, E., and Reichl, U. Comprehensive N-glycosylation analysis of the influenza A virus proteins HA and NA from adherent and suspension MDCK cells. The FEBS Journal.

Pritchard, L.K., Harvey, D.J., Bonomelli, C., Crispin, M., and Doores, K.J. (2015). Cell- and Protein-Directed Glycosylation of Native Cleaved HIV-1 Envelope. Journal of Virology 89, 8932–8944.

Ramakrishnan, B., and Qasba, P.K. (2014). “UDP-Gal: BetaGlcNAc Beta 1,4-Galactosyltransferase, Polypeptide 1 (B4GALT1),” in Handbook of Glycosyltransferases and Related Genes, eds. N. Taniguchi, K. Honke, M. Fukuda, H. Narimatsu, Yamaguchi Y. & T. Angata. (Tokyo: Springer Japan), 51–62.

Rexer, T.F.T., Laaf, D., Gottschalk, J., Frohnmeyer, J., Rapp, E., and Elling, L. (2020a). “Enzymatic synthesis of glycans and glycoconjugates,” in Advances in Glycobiotechnology, eds. E. Rapp & U. Reichl. Springer Nature).

Rexer, T.F.T., Schildbach, A., Klapproth, J., Schierhorn, A., Mahour, R., Pietzsch, M., Rapp, E., and Reichl, U. (2018). One pot synthesis of GDP-mannose by a multi-enzyme cascade for enzymatic assembly of lipid-linked oligosaccharides. Biotechnology and Bioengineering 115, 192–205.

Rexer, T.F.T., Wenzel, L., Hoffmann, M., Tischlik, S., Bergmann, C., Grote, V., Boecker, S., Bettenbrock, K., Schildbach, A., Kottler, R., Mahour, R., Rapp, E., Pietzsch, M., and Reichl, U. (2020b). Synthesis of lipid-linked oligosaccharides by a compartmentalized multi-enzyme cascade for the in vitro N-glycosylation of peptides. Journal of Biotechnology 322, 54–65.

Rödig, J., Rapp, E., Djeljadini, S., Lohr, V., Genzel, Y., Jordan, I., Sandig, V., and Reichl, U. (2011). Impact of Influenza Virus Adaptation Status on HA N-Glycosylation Patterns in Cell Culture-Based Vaccine Production. Journal of Carbohydrate Chemistry 30, 281–290.

Rödig, J.V., Rapp, E., Bohne, J., Kampe, M., Kaffka, H., Bock, A., Genzel, Y., and Reichl, U. (2013). Impact of cultivation conditions on N-glycosylation of influenza virus a hemagglutinin produced in MDCK cell culture. Biotechnology and Bioengineering 110, 1691–1703.

Schön, K., Lepenies, B., and Goyette-Desjardins, G. “Impact of Protein Glycosylation on the Design of Viral Vaccines.” (Berlin, Heidelberg: Springer Berlin Heidelberg), 1–36.

Schwarzer, J., Rapp, E., Hennig, R., Genzel, Y., Jordan, I., Sandig, V., and Reichl, U. (2009). Glycan analysis in cell culture-based influenza vaccine production: Influence of host cell line and virus strain on the glycosylation pattern of viral hemagglutinin. Vaccine 27, 4325–4336.

Selman, M.H.J., Hemayatkar, M., Deelder, A.M., and Wuhrer, M. (2011). Cotton HILIC SPE Microtips for Microscale Purification and Enrichment of Glycans and Glycopeptides. Analytical Chemistry 83, 2492–2499.

Shrotri, M., Swinnen, T., Kampmann, B., and Parker, E.P.K. (2021). An interactive website tracking COVID-19 vaccine development. Lancet Glob Health.

Sommerstein, R., Flatz, L., Remy, M.M., Malinge, P., Magistrelli, G., Fischer, N., Sahin, M., Bergthaler, A., Igonet, S., Ter Meulen, J., Rigo, D., Meda, P., Rabah, N., Coutard, B., Bowden, T.A., Lambert, P.-H., Siegrist, C.-A., and Pinschewer, D.D. (2015). Arenavirus Glycan Shield Promotes Neutralizing Antibody Evasion and Protracted Infection. Plos Pathogens 11.

Stanley, P. (2014). “Mannosyl (alpha-1,3-)-glycoprotein beta-1,2-NAcetylglucosaminyltransferase (MGAT1),” in Handbook of Glycosyltransferases and Related Genes, Second Edition.), 183–194.

Stanley, P., Taniguchi, N., and Aebi, M. (2015). “N-Glycans,” in Essentials of Glycobiology, eds. A. Varki, R.D. Cummings, J.D. Esko, P. Stanley, G.W. Hart, M. Aebi, A.G. Darvill, T. Kinoshita, N.H. Packer, J.H. Prestegard, R.L. Schnaar & P.H. Seeberger. (Cold Spring Harbor (NY): Cold Spring Harbor Laboratory Press), 99–111.

Tsuchiya, S., Aoki, N.P., Shinmachi, D., Matsubara, M., Yamada, I., Aoki-Kinoshita, K.F., and Narimatsu, H. (2017). Implementation of GlycanBuilder to draw a wide variety of ambiguous glycans. Carbohydrate Research 445, 104–116.

Van Landuyt, L., Lonigro, C., Meuris, L., and Callewaert, N. (2019). Customized protein glycosylation to improve biopharmaceutical function and targeting. Current Opinion in Biotechnology 60, 17–28.

Varki, A. (2017). Biological roles of glycans. Glycobiology 27, 3–49.

Walls, A.C., Park, Y.-J., Tortorici, M.A., Wall, A., Mcguire, A.T., and Veesler, D. (2020). Structure, Function, and Antigenicity of the SARS-CoV-2 Spike Glycoprotein. Cell 181, 281–292.

Walls, A.C., Tortorici, M.A., Frenz, B., Snijder, J., Li, W., Rey, F.A., Dimaio, F., Bosch, B.-J., and Veesler, D. (2016). Glycan shield and epitope masking of a coronavirus spike protein observed by cryo-electron microscopy. Nature Structural & Molecular Biology 23, 899–905.

Wan, Y., Shang, J., Graham, R., Baric, R.S., and Li, F. 2020. Receptor Recognition by the Novel Coronavirus from Wuhan: an Analysis Based on Decade-Long Structural Studies of SARS Coronavirus. Journal of Virology [Online], 94. Available: https://jvi.asm.org/content/jvi/94/7/e00127-20.full.pdf.

Watanabe, Y., Allen, J.D., Wrapp, D., Mclellan, J.S., and Crispin, M. (2020). Site-specific glycan analysis of the SARS-CoV-2 spike. Science 369, 330–333.

Watanabe, Y., Bowden, T.A., Wilson, I.A., and Crispin, M. (2019). Exploitation of glycosylation in enveloped virus pathobiology. Biochimica et Biophysica Acta (BBA) - General Subjects 1863, 1480–1497.

Wrapp, D., Wang, N., Corbett, K.S., Goldsmith, J.A., Hsieh, C.L., Abiona, O., Graham, B.S., and Mclellan, J.S. (2020). Cryo-EM structure of the 2019-nCoV spike in the prefusion conformation. Science 367, 1260–1263.

Yang, J., Wang, W., Chen, Z., Lu, S., Yang, F., Bi, Z., Bao, L., Mo, F., Li, X., Huang, Y., Hong, W., Yang, Y., Zhao, Y., Ye, F., Lin, S., Deng, W., Chen, H., Lei, H., Zhang, Z., Luo, M., Gao, H., Zheng, Y., Gong, Y., Jiang, X., Xu, Y., Lv, Q., Li, D., Wang, M., Li, F., Wang, S., Wang, G., Yu, P., Qu, Y., Yang, L., Deng, H., Tong, A., Li, J., Wang, Z., Yang, J., Shen, G., Zhao, Z., Li, Y., Luo, J., Liu, H., Yu, W., Yang, M., Xu, J., Wang, J., Li, H., Wang, H., Kuang, D., Lin, P., Hu, Z., Guo, W., Cheng, W., He, Y., Song, X., Chen, C., Xue, Z., Yao, S., Chen, L., Ma, X., Chen, S., Gou, M., Huang, W., Wang, Y., Fan, C., Tian, Z., Shi, M., Wang, F.-S., Dai, L., Wu, M., Li, G., Wang, G., Peng, Y., Qian, Z., Huang, C., Lau, J.Y.-N., Yang, Z., Wei, Y., Cen, X., Peng, X., Qin, C., Zhang, K., Lu, G., and Wei, X. (2020). A vaccine targeting the RBD of the S protein of SARS-CoV-2 induces protective immunity. Nature 586, 572–577.

Zhou, P., Yang, X.-L., Wang, X.-G., Hu, B., Zhang, L., Zhang, W., Si, H.-R., Zhu, Y., Li, B., Huang, C.-L., Chen, H.-D., Chen, J., Luo, Y., Guo, H., Jiang, R.-D., Liu, M.-Q., Chen, Y., Shen, X.-R., Wang, X., Zheng, X.-S., Zhao, K., Chen, Q.-J., Deng, F., Liu, L.-L., Yan, B., Zhan, F.-X., Wang, Y.-Y., Xiao, G.-F., and Shi, Z.-L. (2020). A pneumonia outbreak associated with a new coronavirus of probable bat origin. Nature 579, 270–273.

