## Supplementary Information for "Cell-free glycoengineering of the recombinant SARS-CoV-2 spike glycoprotein"

### *Supplementary Material*

#### 1 Supplementary Data

##### 1.1 Additional information on enzymes

For gene expression *E. coli* strains were typically grown to an $OD_{600}$ of 0.6-0.8 in LB/TB media at 30°C followed by induction using 0.4 mM IPTG and a reduction of the cultivation temperature to 16°C. Plasmids with the gene insert were purchased from BioCat (Heidelberg, Germany).

##### 1.2 Results

The recombinant enzymes were analyzed by SDS-PAGE after IMAC purification confirming production of all His-tagged variants (Supplementary Figure 1).

### 2 Supplementary Figures and Tables

#### 2.1 Supplementary Figures

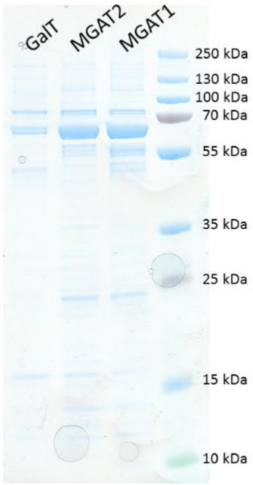


**Supplementary Figure 1.** SDS-PAGE (12% Bis-Tris) of 4 µg GalT, MGAT2 and MGAT1 each. Theoretical protein masses are: MGAT1ΔTM = 50.9 kDa, MGAT2ΔTM = 54.4 kDa , and GalTΔTM = 45.5 kDa .

#### 2.2 Supplementary Tables

**Supplementary Table 1.** List of chemicals. Suppliers: AppliChem (Darmstadt, Germany), Applied Biosystems (Waltham, USA), Carl Roth (Karlsruhe, Germany), glyXera (Magdeburg, Germany), Merck (Darmstadt, Germany), Sigma Aldrich (St. Louis, USA) [now Merck], Thermphos International (Wittenberg, Germany), Thermo Scientific (Waltham, USA):

| **Chemical** | **Supplier** | **Product number** | **Purity** |
|---|---|---|---|
| glyXprep16™ kit | glyXera | KIT001-16S | - |
| 8-aminopyrene-1,3,6-trisulfonic acid (ATPS) | Sigma | 09341 | > 96% |
| Acetonitril | Thermo Scientific | A955 | Optima™ "LC/MS grade" |
| BCA assay kit | Thermo Scientific | 23227 | - |
| Glycerol | Carl Roth | 3783.1 | >99.5% |
| HCl | Carl Roth | 4025 | 37 % |
| HEPES | Carl Roth | 9105.3 | ≥99.5 % |
| HiDi™-formamide | Applied Biosystems | 4311320 | - |
| Kanamycin sulfate | Merck | 10106801001 | - |
| LIZ™ | Applied Biosystems | 4322679 | - |
| Imidazole | Carl Roth | 3899.4 | - |
| Isopropyl β-D-1-thiogalactopyranoside (IPTG) | AppliChem | A1008,0025 | - |
| $MnCl_2$ | Merck | 1.05934.0100 | - |
| NaCl | Carl Roth | P029.3 | ≥99 % |
| Trifluoroacetic acid | Merck | 302031 | >99% |
| Tris(hydroxymethyl)-aminomethan-buffer (TRIS) | AppliChem | A2264 | > 99.9%) |
| Tryptone | Carl Roth | 8952.2 | - |

| | | | |
|---|---|---|---|
| UDP-GlcNAc | Carbosynth | MU07955 | - |
| UDP-galactose | Carbosynth | MU06699 | - |
| Yeast extract | Carl Roth | 2363.2 | |

**Supplementary Table 2.** Genes, vectors and strains used for the synthesis of recombinant glycosyltransferases.

| **Enzymes** | **Uniprot ID** | ***E. coli* strain** | **Plasmid** |
|---|---|---|---|
| MGAT1 | P26572 | BL21(DE3) | pET-28a(+) |
| MGAT2 | Q10469 | SHuffle® T7 *lysY* | pET-28b(+) |
| GalT | P15291 | BL21(DE3) | pET-28a(+) |
